# Deltaviruses spread through a viral Trojan Horse

**DOI:** 10.1101/2025.05.09.653040

**Authors:** Joe McKellar, Aurélien Fouillen, Sébastien Lyonnais, Florian Seigneuret, Marie-Pierre Blanchard, Antonio Trullo, Lorena Kumarasinghe, Sylvain De Rossi, Roni Sleiman, Solange Desagher, Raphael Gaudin, Hugues de Rocquigny, Sébastien Granier, Jussi Hepojoki, Karim Majzoub

## Abstract

Hepatitis D-like viral satellites, known as deltaviruses, have been recently discovered in a wide range of animals. These viruses are thought to expropriate glycoproteins from helper viruses to form infectious particles. Here, we challenge this paradigm and demonstrate that deltaviruses are packaged within helper virus particles, using them as viral Trojan Horses for cell entry. By leveraging orthogonal electronic and photonic super-resolution microscopy, we visualize deltaviruses enclosed within virions from rhabdo-, herpes-, and arenavirus families. We show that this conserved hitchhiking mechanism ensures concomitant deltavirus-helper virus spread, which advantages the dissemination of deltaviruses, broadens their host range and expands their tropism. Our findings reveal a previously unrecognized mode of viral transmission, providing a framework to investigate overlooked deltavirus infections outside of the human liver.

## Introduction

Viral satellites have been found across many eukaryotic species such as plants and insects ^1^. Until very recently the only two known viral satellites known to infect vertebrate animals, and more particularly humans, were 1-members of the adeno-associated virus family (AAVs) (*Parvoviridae*) and 2-Hepatitis D virus (HDV). HDV depends on the Hepadnavirus Hepatitis B virus (HBV) surface glycoproteins for infection ^2^, and their co-infection causes the most severe form of viral hepatitis. A few years ago, the simultaneous discovery of HDV-*like* sequences in many animal hosts such as rodents, bats, birds, snakes, frogs and even insects ^3–10^ revealed that HDV belongs to a large family of negative-sense, circular viral RNA satellites that are widespread across the animal kingdom, now called *Kolmioviridae* ^11^. Importantly, unlike HDV, these other animal deltaviruses were not only found in the liver of infected hosts but in several other organs including gastrointestinal tract, lung, blood, spleen and brain tissues, with no evidence of HBV or other Hepadnavirus co-infection ^3,4,6^. Initial reports indicate that the Snake deltavirus (SDeV; also called Swiss snake colony virus 1), discovered in boa constrictors co-infected with reptarenaviruses, uses these viruses as helpers ^4,12^. Apart from SDeV, the identity of helper viruses associating with the vast majority of these newly discovered RNA satellites remains unknown. Helper virus glycoproteins have been shown to play a role in deltavirus infectivity ^13,12,14^, however the nature of deltavirus associations with their helper virus counterparts is unknown. A better understanding of the rules governing how deltaviruses co-opt helper viruses to spread is crucial because successful associations would influence deltavirus species range and tissue tropism ^7^. In this study, we show that deltaviruses spread through a viral Trojan Horse, after being packaged within their helper virus particles. We also provide evidence that this novel mode of viral transmission ensures concomitant deltavirus-helper virus spread, dictating the tissue- and species-tropism of these circular RNA viral satellites.

## Results

### VSV virions are morphologically modified upon superinfection of deltavirus-replicating cells

We have previously shown that the neotropical Tome’s spiny rat (*Proechimys semispinosus*) rodent deltavirus (RDeV) ^6^ is able to replicate in a wide range of animal cells and can be packaged with the vesicular stomatitis virus (VSV) glycoprotein (VSV-G) to form infectious particles ^14^. To explore this in a more physiological context, we superinfected a mouse fibroblast cell-line persistently replicating RDeV (NIH3T3-RDeV) ^14^ with a full, replication competent VSV and monitored deltavirus virion production (Fig. 1A). We detected by immunoblot rodent delta antigen (RDAg) in supernatants of VSV-superinfected NIH3T3-RDeV cells (Fig. 1B) and observed by RT-qPCR a ∼25-fold increase in supernatant RDeV RNA levels compared to those of non-superinfected cells (Fig. 1C), with no impact of RDeV on VSV RNA levels (Fig. 1D). This indicated that while some residual RDeV RNA is secreted from cells at baseline levels, VSV superinfection greatly stimulated both RDAg and RDeV RNA secretion. To explore the nature of secreted RDeV virions, we performed Negative Stain Transmission Electron Microscopy (NS-TEM) on these supernatants (Fig. 1E). VSV-superinfected NIH3T3-RDeV supernatants contained characteristic bullet-shaped VSV virions ^15^ (Fig. 1E, inset 1) and smaller spherical particles (av. 43.28 nm in diameter) (Fig. 1E, inset 2, 1F), supposedly RDeV virions, as absent in control conditions (Fig. S1A), and, while slightly larger, are consistent with reported HDV particle dimensions ^13,16^. Surprisingly, we identified a third class of particles, being VSV virions physically associated with RDeV particles (Fig. 1E, inset 3). 13% of observed VSV virions were associated with these smaller particles (Fig. 1G). While 13% is probably an underestimation—false negatives in the classification as well as hidden surface areas on particles—it nevertheless outnumbered virions associated with supernatant debris in RDeV-free conditions (2%) (Fig. 1G). Furthermore, these VSV-associated particles had a smaller average size (av. 35.5 nm) compared to the free RDeV particles, raising the possibility of them being extensions of VSV virions, with the deltavirus ribonucleoprotein (vRNP) being contained within (Fig. 1F). Of note, these VSV-associated particles did not localize preferentially to a specific region of VSV (Fig. 1E, H, S1B). Also of note, some VSV virions were associated with more than one (up to 4) of such particles, exclusively in the presence of RDeV (Fig. S1B). We were also able to visualize similar VSV-associated particles in supernatants produced from NIH3T3 cells persistently replicating the woodchuck (*Marmota monax*) deltavirus (mmDeV) ^8,14^ (NIH3T3-mmDeV) with comparable frequency (Fig. 1G-H), suggesting that these particles could be a general feature of deltaviruses. Next, to rule-out chemical fixation artifacts (required prior to BSL-3 exit) as the origin of these VSV-RDeV particles, we performed Bio-Atomic Force Microscopy (Bio-AFM) on non-fixed VSV-superinfected NIH3T3-RDeV supernatants within the BSL-3 environment. Bio-AFM assesses the topological properties of native infectious particles at single virion resolution ^17^. Similar to NS-TEM, Bio-AFM detected free VSV particles, free RDeV particles as well as VSV-associated RDeV particles exclusively from NIH3T3-RDeV supernatants (Fig. 1I-J, S1C-G). The dimensions of VSV-associated RDeV particles and free RDeV particles are consistent with those measured by NS-TEM (Fig. 1F, 1K), confirming that VSV-associated RDeV particles were smaller in size than free RDeV particles (av. 27 nm vs 36 nm, respectively). Of note, a cantilever-induced compression of free and VSV-associated RDeV particles is observed (Fig. 1K), as expected ^18^, hence the slightly smaller size compared to NS-TEM (Fig. 1F). Conversely, VSV virions were not deformed (av. 67 nm vs 70 nm for Cryo-EM ^19^), suggesting that deformability of deltaviruses under the Bio-AFM is likely due to the lack of a matrix protein associated with their membrane. Also, in agreement with NS-TEM, we found VSV decorated with more than one RDeV particle (Fig S1G). Therefore, VSV-associated deltaviruses occur natively and are not fixation artefacts. Altogether, our data suggests that deltaviruses may be associated with or contained within VSV virions.

**Fig. 1.**
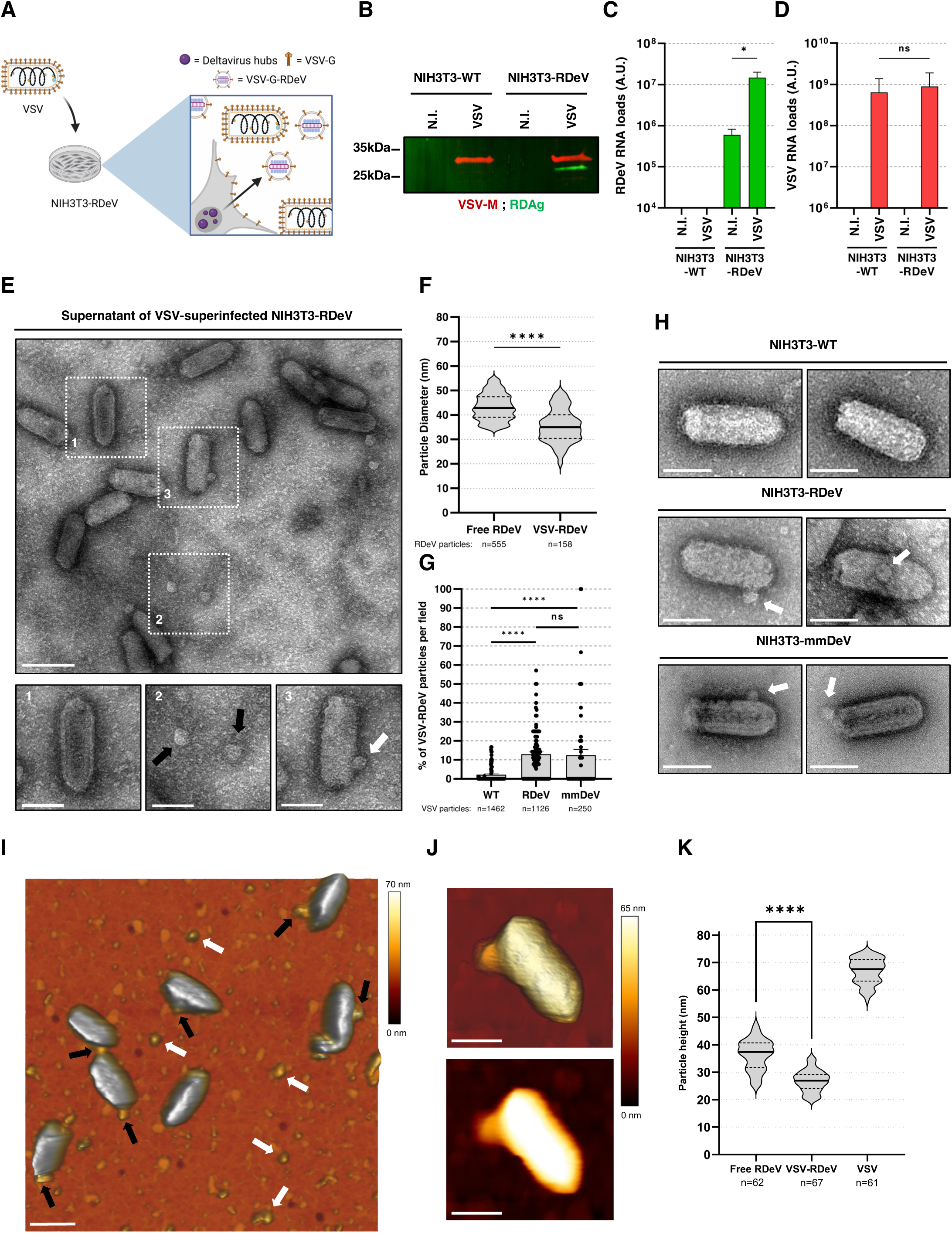
A subset of VSV virions is morphologically modified upon superinfection of deltavirus-replicating cells. (**A**) Schematic representation of the superinfection of rodent deltavirus (RDeV)-replicating cells by VSV. (**B**) Immunoblot of equal volumes of supernatants of non-infected (N.I.) or VSV-infected NIH3T3-WT or -superinfected NIH3T3-RDeV cells 16 h post-infection. VSV-M is represented in red and RDAg in green. (**C**) RT-qPCR of RDeV RNA loads from equivalent volumes of supernatants of B. (**D**) RT-qPCR of VSV RNA loads from equivalent volumes of supernatants of B. (**E**) Representative NS-TEM image of the supernatants of B. Inset 1, 2 and 3 show free VSV particles, free RDeV particles (black arrows) and VSV-associated RDeV particles (white arrows), respectively. Scale bars: 200 nm, insets: 100 nm. (**F**) Quantification of the average size of free or VSV-associated RDeV particles. A total of 555 free RDeV particles and 158 VSV-associated RDeV particles were analyzed in total. (**G**) Quantification of the percentage of RDeV-associated VSV virions per field. A total of 100, 99 and 58 fields, corresponding to 1462, 1126 and 250 VSV particles, were analyzed for deltavirus-free, RDeV and mmDeV conditions, respectively. (**H**) Representative NS-TEM images of VSV particles in WT, RDeV or mmDeV conditions, showing VSV-associated deltavirus particles (white arrow). Scale bars: 100 nm. (**I**) AFM image of native VSV particles produced from NIH3T3-RDeV cells, the white and black arrows point to free RDeV and VSV-associated RDeV particles, respectively. Scale bar: 200 nm (**J**) 3D projection and AFM topographic image of a RDeV-associated VSV virion. Scale bars: 100 nm. (**K**) Quantification of the average particle height of free RDeV particles (n=62), VSV-associated RDeV (n=67) or VSV virions (n=61). Experiments are representative of 2 independent repeats, performed in technical duplicates for C-D. Unpaired T-tests were used to evaluate statistical significance.

### Deltaviruses are packaged inside VSV virions using them as a viral Trojan Horse for cell entry

Next, to ascertain the identity of the previously observed VSV-associated particles, we used specific VSV and deltavirus antibodies ^14^ to stain VSV-superinfected NIH3T3-RDeV cell secretions and analyzed virions by Stimulated Emission Depletion (STED) and Airyscan super-resolution microscopy (Fig. 2A, S2A). As expected, both techniques revealed the presence of elongated VSV particles, smaller free RDeV particles and confirmed that some VSV particles stained locally positive for RDAg, at the periphery of VSV virions, absent in control conditions (Fig. 2A, S2A). Quantification of the STED images showed that ∼25%—over background—of VSV particles co-stained for RDAg, with ∼18% of RDeV particles being associated to VSV (Fig. 2B-C). We further orthogonally validated these results by immunogold-EM, which showed a local accumulation of RDAg at the periphery of ∼20%—over background—of VSV virions (Fig. 2D-E). Similar results were also obtained with VSV-associated mmDeV particles by both STED (Fig. 2B-C, S2B) and immunogold (Fig. 2D-E, S2C). These approaches confirmed that the VSV-associated structures observed earlier in NS-TEM and Bio-AFM (Fig. 1) are indeed deltaviruses.

**Figure 2.**
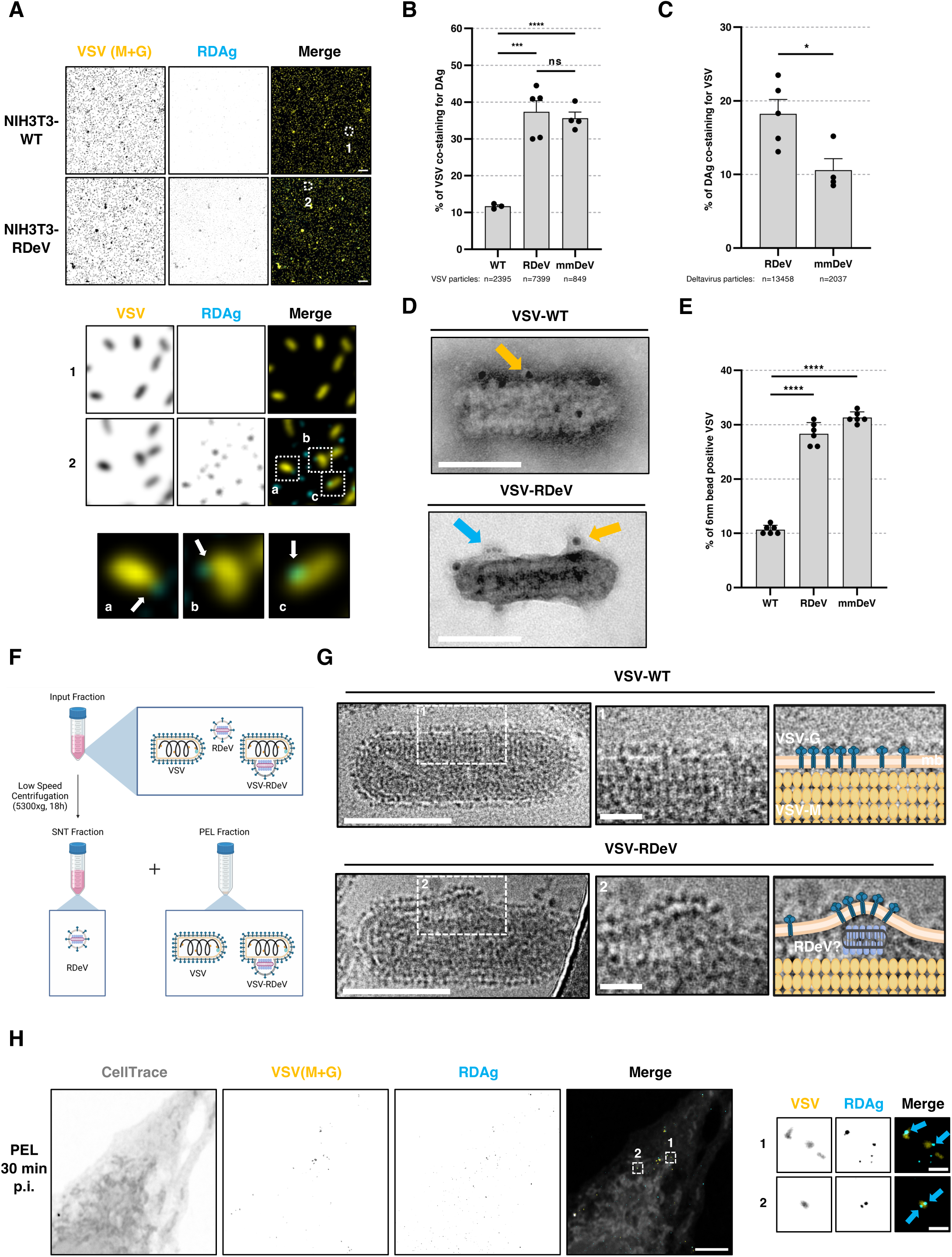
Deltaviruses are packaged inside VSV virions and use them as a viral Trojan Horse for cell entry. (**A**) Representative STED images of VSV-infected NIH3T3-WT or -superinfected NIH3T3-RDeV supernatants. VSV (M+G) is represented in yellow and RDAg in cyan. Single channels are shown in inverted grey. White arrows show RDeV particles associated to VSV particles. Scale bar: 1 µm. (**B**) Quantification of the percentage of VSV co-staining with DAg from the STED images. A total of 2395, 7399 or 849 VSV particles were quantified for WT, RDeV and mmDeV conditions, respectively. (**C**) Quantification of the percentage of RDeV or mmDeV DAg co-staining with VSV from the STED images. A total of 13458 RDeV or 2037 mmDeV particles were quantified. (**D**) Representative NS-TEM images of immunogold labeling of VSV-G (10 nm gold beads; yellow arrows) and RDAg (6 nm gold beads; blue arrow) from supernatants of VSV-infected NIH3T3-WT or -superinfected NIH3T3-RDeV cells. Scale bar: 100 nm. (**E**) Quantification of the immunogold experiments showing percentage of VSV particles positive for both 10 nm beads (VSV-G) and 6 nm beads (RDAg). (**F**) Schematic representation of the Low-Speed Centrifugation (LSC) pipeline (PEL for pellet fraction and SNT for supernatant fraction) (**G**) Representative cryo-EM micrographs of VSV-WT or VSV-RDeV virions. Inset 2 zooms on the local membrane elevation caused by a density probably being the RDeV vRNP. A total of 44 of these events were seen over 137 fields. Scheme (right panel) represents VSV membrane (mb) in orange, VSV-M in yellow, VSV-G in dark blue and the RDeV vRNP (RNA in gray and RDAg in light blue), superimposed on insets 1 and 2. Scale bars: 100 nm, inset: 20 nm. (**H**) STED image of a Vero cell pre-stained with Celltrace-CFSE and infected with VSV-RDeV LSC PEL fraction for 30 min. VSV (M+G) proteins are stained in yellow and RDAg in cyan. Insets show VSV-RDeV virions (blue arrows) localizing inside of the cell. Scale bar: 5 µm; inset: 500 nm. Experiments are representative of 2 independent repeats. Unpaired T-tests were used to evaluate statistical significance.

We then sought to gain insight into the nature of the observed VSV-deltavirus associations. First, we took advantage of a VSVΔG replication-competent virus ^20^, that produces VSV-G devoid virions ^21^, to determine if VSV-G is essential for VSV-RDeV association. NS-TEM revealed the presence of VSVΔG-RDeV particles (Fig. S2D), indicating that VSV-RDeV association is independent of the fusogenic property of VSV-G ^22^. This suggests that VSV-RDeV do not associate through VSV-G fusion in the extracellular milieu but rather intracellularly during VSV budding, as a single entity. To determine if a continuous viral membrane surrounds these VSV-RDeV particles, we set up a Cryo-EM approach. Low Speed Centrifugation (LSC) ^23^ of viral supernatants separated free deltavirus particles in the LSC supernatant (SNT) from VSV-RDeV particles in the LSC pellet (PEL) fractions (Fig. 2F, S2E-F). Cryo-EM of the NIH3T3-RDeV LSC PEL fraction revealed the existence of a population of VSV particles that presented a local density between the matrix layer and the virion membrane, absent in control conditions (Fig. 2G). This density, locally elevating the virion membrane, likely corresponds to the RDeV vRNP, nevertheless future studies should aim to model the 3-dimensional organization of this vRNP. This data therefore implies that RDeV is contained within the VSV virion, forming a single viral unit. This unique virion organization could benefit deltavirus propagation, allowing them to hitchhike within helper virus particles to concomitantly infect the same host cell. We propose to name this propagation model viral Trojan Horse spread.

To confirm such a model, we probed early viral entry events, infecting Vero cells for 30 min with VSV-superinfected NIH3T3-RDeV LSC input, SNT or PEL fractions, followed by STED microscopy. This revealed free RDeV particle entry from the Input and SNT fractions (Fig. S2G) as well as VSV-RDeV particle entry from the PEL and Input fractions (Fig. 2H, S2G), where VSV-RDeV particles were seen among intracellular membranes. This confirms the model by which deltaviruses are able to hitch-hike with VSV during a superinfection, as a viral Trojan Horse, ensuring concomitant transmission and entry into naïve cells.

### Viral Trojan Horse spread is indispensable for deltavirus association with herpesvirus

Following VSV-RDeV entry, a limitation to follow deltavirus replication over time is the cytolytic capacity of VSV. Indeed, VSV completes a single round infection in around 6 h ^24^ and is cytolytic rapidly after a few infection cycles ^25^, whereas the detection of replicating RDeV requires several days of infection ^6,14^. Consequently, we explored another model virus for its viral Trojan Horse potential, the dsDNA Herpes Simplex Virus 1 (HSV-1). Importantly, HSV-1 can enter into latency in neuronal tissues ^26^, induces cytopathic effects in 2-3 days ^27^ and possesses a large 200 nm virion ^28^ that could accommodate the deltavirus vRNP, making it an ideal Trojan Horse helper virus in theory.

Superinfection of NIH3T3-RDeV with HSV-1 KOS-64-GFP (hereafter, HSV-1) caused an ∼100-fold increase in extracellular secretion of RDeV genome compared to baseline levels (Fig. 3A). Visualization of HSV-1 by NS-TEM was unsuccessful and our Cryo-EM setup did not allow us to visualize the inside of HSV-1 virions (Fig. S3A), so we therefore turned to specific antibody staining followed by STED and Airyscan microscopy (Fig. 3B, S3B). Secretions from HSV-1 superinfected NIH3T3-RDeV cells revealed the presence of free HSV-1 particles, free RDeV particles and HSV-1 particles that co-stained locally for RDAg (Fig. 3B) at a frequency of ∼40%— above background— (Fig. 3C), suggesting that RDeV is contained within HSV-1 virions. Furthermore, some HSV-1 particles stained positive for multiple instances of RDAg (Fig. 3B, inset 2). To functionally test if HSV-1-RDeV virions benefit deltavirus infectivity we separated by low-speed ultracentrifugation free RDeV particles from HSV-1 and HSV-1-RDeV particles (Fig. 3D). As expected, the majority of RDeV RNA, RDAg and particles remained in the SNT fraction, whereas HSV-1 particles were concentrated in the pellet (Fig. 3E, S3C-D). These different fractions were then used to infect Vero cells in the presence of Acyclovir, an HSV-1 inhibitor, to monitor RDeV replication (Fig. 3F). Surprisingly, SNT fractions, containing free RDeV particles, were unable to initiate any RDeV infection, whereas RDeV-positive Vero cells were observed with PEL fraction infections, where HSV-1-RDeV particles resided (Fig. 3F). Some cells, where Acyclovir did not fully inhibit HSV-1, were double-positive for HSV-1 and RDeV, showing concomitant entry and replication of both viruses (Fig. 3G). Furthermore, we could observe nuclear deltavirus hubs, a hallmark of deltavirus replication and accumulation ^14^ (Fig. 3G). Importantly, the fact that free RDeV particles were not infectious suggests the mandatory reliance of RDeV on HSV-1, as a viral Trojan Horse, for infection. To confirm this, we overexpressed HSV-1 glycoproteins (gB, gD, gH and gL) ^29^ or VSV-G in NIH3T3-RDeV cells and collected cell secretions to infect Vero cells (Fig. S3F). Unlike with VSV-G ^14^, HSV-1 glycoproteins were unable to pseudotype infectious RDeV particles (Fig. S3E). Interestingly, the glycoproteins of another herpesvirus, Human cytomegalovirus (HCMV; gH, gL and gO) ^30^, were also unable to pseudotype infectious RDeV particles (Fig. S3E). This data confirms the mandatory reliance of RDeV on the viral Trojan Horse spread, in the case of herpesviruses, for infectivity.

**Figure 3.**
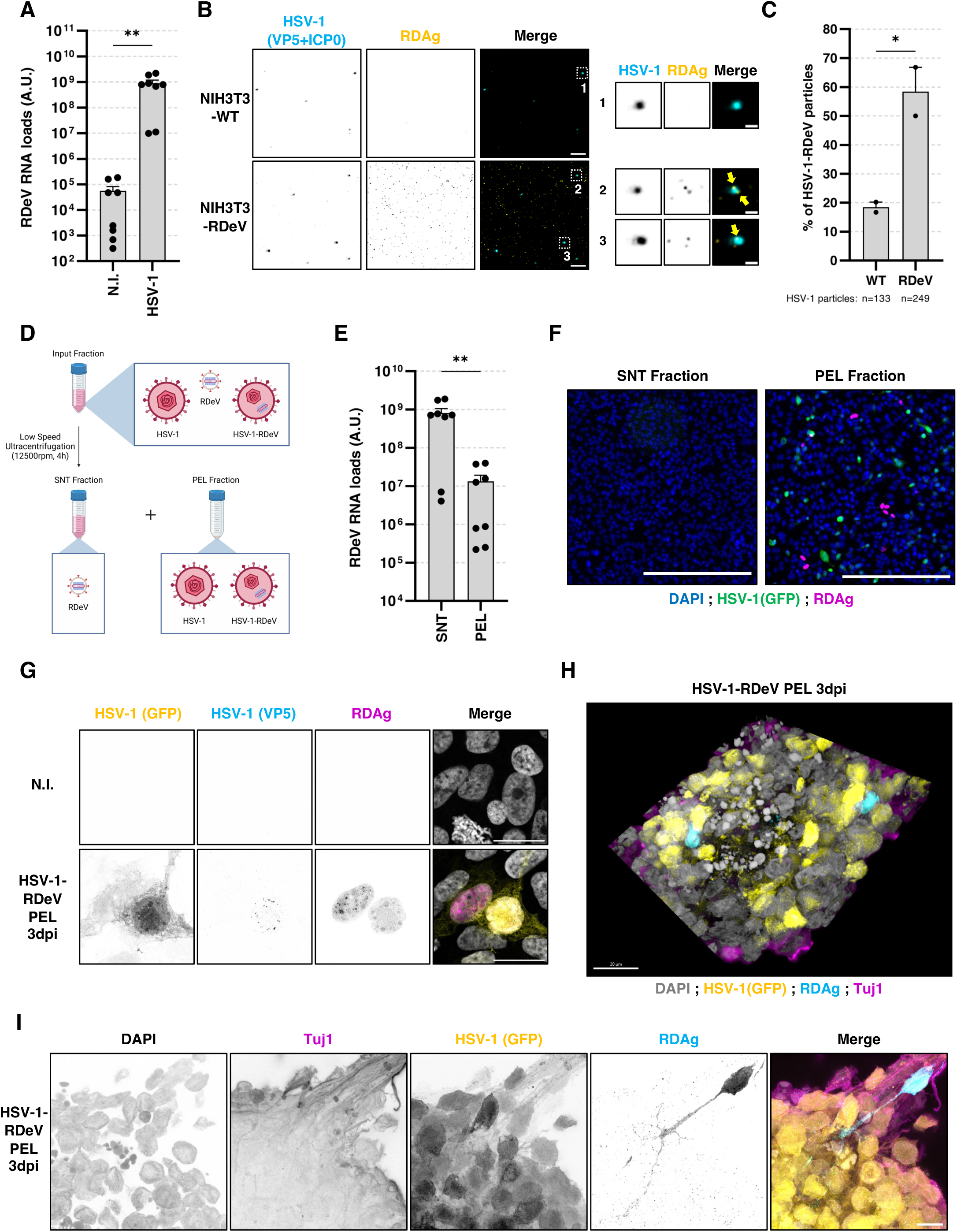
Viral Trojan Horse spread is indispensable for deltavirus association with HSV-1 for productive infection. (**A**) RDeV RT-qPCR of volume equivalent supernatants of non-infected (N.I.) or HSV-1 superinfected NIH3T3-RDeV 72 h post-infection. (**B**) Representative STED images of HSV-1-infected NIH3T3-WT or -superinfected NIH3T3-RDeV supernatants. HSV-1 (VP5+ICP0) is represented in cyan and RDAg in yellow. Single channels are shown in inverted grey. Yellow arrows point to RDAg puncta associated with HSV-1 particles. Scale bars: 1 µm, inset: 200 nm. (**C**) Quantification of the percentage of HSV-1 virions co-staining positive for RDAg in the supernatant of HSV-1-infected NIH3T3-WT or superinfected-RDeV from STED imaging. 133 virions were analyzed for the WT condition and 239 for the RDeV condition. (**D**) Schematic representation of the low-speed ultracentrifugation strategy of HSV-1- and RDeV-containing supernatants (PEL for pellet fraction and SNT for supernatant fraction). (**E**) RDeV RT-qPCR of volume equivalent SNT and PEL fractions of C. (**F**) Representative epifluorescence images of Vero cells infected with SNT or PEL fractions 2 days post-infection in the presence of 50 µM HSV-1 inhibitor Acyclovir to limit cell death. Nuclei are shown in blue, HSV-1 (GFP) in green and RDAg in magenta. Scale bar: 200 µm. (**G**) Representative Airyscan images of Vero cells infected with the HSV-1-RDeV PEL fraction 3 days post-infection in the presence of 50 µM Acyclovir. Nuclei are shown in grey, HSV-1 (GFP) in yellow, HSV-1 (VP5) in cyan and RDAg in magenta. Single channels are shown in inverted greys. Scale bar: 20 µm. (**H**) Representative confocal z-stack 3D-rendering of a field of LUHMES-derived neurons infected with HSV-1-RDeV PEL fraction 3 days post-infection in the absence Acyclovir. Nuclei are shown in grey, HSV-1 (GFP) in yellow, RDAg in cyan and Tuj1 (neuronal marker) in magenta. Scale bar: 20 µm. Corresponding animated 3D-rendering of this z-stack can be found in Movie S1. (**I**) Airyscan z-stack maximum projection of a field of LUHMES-derived neurons infected with HSV-1-RDeV PEL fraction 3 days post-infection in the absence Acyclovir. Nuclei are shown in grey, HSV-1 (GFP) in yellow, RDAg in cyan and Tuj1 in magenta. Single channels are shown in inverted grays. Scale bar: 10 µm. Corresponding animated 3D-rendering of this z-stack can be found in Movie S2. Experiments are representative of 2 (B, C, F-G), 3 (H-I) or 4 (A, E) independent repeats. Unpaired T-tests were used to evaluate statistical significance.

RDeV was originally found in the blood, kidney, heart, small intestine and lungs of infected spiny rats ^6^ and is able to replicate in many different cell lines ^14^. The natural tropism of HSV-1 being human neurons, we were able to show that in the absence of Acyclovir, HSV-1-RDeV from the PEL fraction can also infect post-mitotic neurons derived from the human neuronal precursor cell line LUHMES (Fig. 3H-I, Movie S1-S2). In these cells, RDeV presence was always associated to HSV-1 positivity (Fig. 3H-I, Movie S1-S2), and in some cases colonized the neuron prolongations as well as the nucleus (Fig. 3I, Movie S2). Furthermore, as expected (Fig. 3E), the SNT fraction remained non-infectious (Fig. S3F, Movie S3), reconfirming that HSV-1-RDeV Trojan Horse particles are the only effective way for RDeV to spread with HSV-1. Furthermore, the ability of RDeV to infect human neurons highlights the tropism and host-shifting capacity that viral Trojan Horse transmission enables when deltaviruses are packaged within herpes virions.

### Snake deltavirus uses its natural helper arenavirus as a viral Trojan horse

Although our experimental data clearly shows that deltavirus association with rhabdo- and herpesviruses is possible, these associations have not yet been observed in naturally infected animals. However, SDeV has been found associated with arenaviruses in boa constrictor snakes and was originally isolated from the brain of an infected boa constrictor along with these helper viruses ^4,12^. We therefore turned towards SDeV and explored the nature of its association with the reptarenavirus University of Giessen virus-1 (UGV-1) ^31^, as a viral Trojan Horse.

We superinfected I/1Ki boa constrictor cells ^4^ replicating SDeV (I/1Ki-SDeV) with UGV-1 and recovered the supernatant 12 days post-infection. We saw a modest but significant increase in SDeV RNA secretion upon superinfection (Fig. 4A), with no impact of SDeV on UGV-1 viral RNA levels at the time of collection (Fig. 4B). Separation of infectious particles by centrifugation (Fig. 4C) showed that the majority of the SDeV RNA and snake delta antigen (SDAg) remained in the SNT fraction, with only a minority pelleting with UGV-1 (Fig. 4D-F). We next explored the nature of UGV-1 particles in absence or presence of SDeV by NS-TEM. Morphologically, UGV-1 particles produced in I/1Ki-WT revealed a pleiomorphic virion size, as expected (Fig. 4G) ^32^. Remarkably, ∼12% of UGV-1 particles produced in I/1Ki-SDeV cells contained an additional density within the virion (Fig. 4H-I), supposedly being the SDeV vRNP, as never seen under control conditions (Fig. 4G, I). STED imaging using reptarenavirus and SDeV specific antibodies, revealed the presence of SDeV-positive UGV-1 particles (Fig. 4J), with a frequency of ∼19%— over background—(Fig. 4K), confirming that the observed density in NS-TEM corresponds to SDeV vRNP within UGV-1 particles. Finally, monitoring infection of free SDeV from the SNT fraction versus UGV-1-SDeV from the PEL fraction showed an ∼300-fold higher infectivity of the PEL versus the SNT fraction (Fig. 4L). This suggests that UGV-1-SDeV virions favor SDeV infectivity compared to SDeV free particles decorated with UGV-1 glycoproteins. Our data therefore show that SDeV is able to use UGV-1 as a viral Trojan Horse favoring SDeV infectivity compared to glycoprotein-decorated free SDeV particles. This data therefore highlights that viral Trojan Horse spread occurs between SDeV and its natural helper virus, solidifying this mode of viral transmission as a generality of deltavirus association with their helper viruses (Fig. S4).

**Figure 4.**
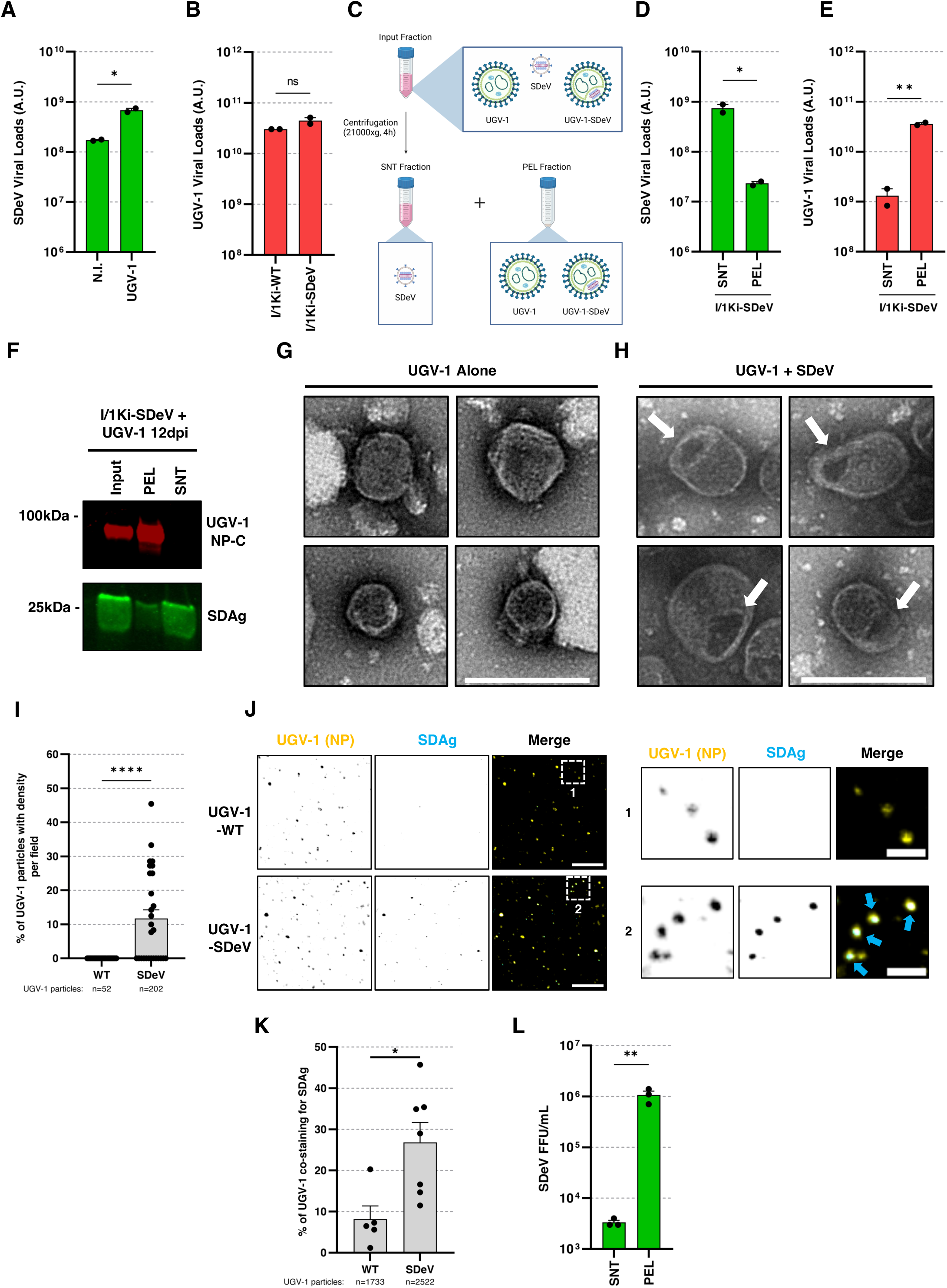
Snake deltavirus uses UGV-1 as a viral Trojan horse. (**A**) SDeV RT-qPCR of volume equivalent supernatants of non-infected or UGV-1-superinfected I/1Ki-SDeV cells 12 days post-infection. (**B**) UGV-1 RT-qPCR of volume equivalent supernatants of UGV-1 infected I/1Ki-WT or superinfected I/1Ki-SDeV cells 12 days post-infection. (**C**) Schematic representation of the centrifugation strategy of UGV-1- and SDeV-containing supernatants (PEL for pellet fraction and SNT for supernatant fraction). (**D**) SDeV RT-qPCR of volume equivalent SNT and PEL fractions from C. (**E**) UGV-1 RT-qPCR of volume equivalent supernatants of C. (**F**) Representative immunoblot of volume equivalent centrifugation fractions from C. UGV-1 NP-C is shown in red and SDAg in green. **G.** NS-TEM images of UGV-1 particles from I/1Ki-WT. (**H**) NS-TEM images of UGV-1 particles from I/1Ki-SDeV. White arrows show a density inside of UGV-1 particles, found uniquely in the I/1Ki-SDeV supernatants. Scale bar: 100 nm. (**I**) Quantification of the percentage of UGV-1 particles containing a density per field. 27 and 28 fields, with a total of 52 and 202 UGV-1 particles, were analyzed for I/1Ki-WT and -SDeV conditions respectively. (**J**) Representative STED images of UGV-1-infected I/1Ki-WT or - superinfected I/1Ki-SDeV PEL fractions. Cyan arrows point to SDAg inside of UGV-1 particles. Scale bar: 1 µm; inset: 500 nm. (**K**) Quantification of the percentage of UGV-1 particles co-staining for SDAg from the STED images. A total of 1733 and 2522 UGV-1 particles were analyzed for WT and SDeV conditions respectively. (**L**) SDeV FFU/mL of naïve I/1Ki cells from the SNT and PEL fractions 3 days post-infection. Experiments are representative of 2 independent repeats Unpaired T-tests were used to evaluate statistical significance.

## Discussion

Herein, our study demonstrates that animal deltavirus vRNPs can be packaged within helper virus virions. This property allows them to hitchhike, using helper virions as Trojan Horse vehicles to facilitate their spread and entry into new host cells, simultaneously ensuring co-infection with the helper virus. Since entry of enveloped viruses is a key determinant of the species barrier ^33^, this unconventional Trojan Horse mechanism may contribute to the host-shifting capacity of deltaviruses ^7^. Furthermore, given their ability to replicate in diverse cell types ^14^ this transmission strategy likely influences their tissue tropism. This may explain why deltavirus genomes have been detected across multiple infected animal tissues ^3,4,6^. Interestingly, while deltaviruses can associate with envelope glycoproteins of VSV and UGV-1 to form independent infectious particles ^12,14^—without requiring enclosure within helper virus particles—our data indicates that viral Trojan Horse propagation achieves higher deltavirus infection rates (Fig. 3F, 4L). Furthermore, in the case of HSV-1, the Trojan Horse mechanism is the only means by which deltaviruses can be infectious (Fig. 3). A possible explanation for this discrepancy is the differential expression of envelope glycoproteins from different viral families and their distinct budding sites. Both VSV and UGV-1 express their glycoproteins at the plasma membrane, where they bud ^19,34^, potentially allowing deltavirus vRNPs to be co-packaged. In contrast, HSV-1 buds intracellularly in the Golgi apparatus ^35^ and in the absence of HSV-1 tegumentation ^36^, deltaviruses might never be able to associate with HSV-1 glycoproteins (Fig. S4). Varying rates of deltavirus vRNP incorporation into heterologous viral particles further suggest that this process may be stochastic and inefficient, particularly given that only a minority of helper virions contain deltavirus vRNPs (25% for VSV, 40% for HSV-1, 19% for UGV-1, Fig. 2B, S3C, 4K). It is therefore conceivable that deltaviruses have evolved an inefficient, yet highly generalist, co-packaging strategy enabling them to associate with a wide range of viral families with diverse assembly and budding mechanisms. Despite its inefficiency, viral Trojan Horse spread may still be more advantageous for deltaviruses than propagating as free particles bearing heterologous viral glycoproteins without encoding them. The Trojan Horse mechanism ensures that deltaviruses enter host cells simultaneously with the genetic material encoding their glycoproteins and vehicle, thereby enhancing the likelihood of remaining associated with its helper virus in subsequent rounds of infection.

Importantly, while physical associations between satellite and helper viruses have been observed in other domains of life such as the satellite-helper phage system infecting *Streptomyces* species ^37^ and virophages interacting with giant viruses in amoebae ^38^—our study presents the first known example of a viral satellite contained within an animal virus particle. Unlike bacterial phages and giant amoebal viruses, the association described here likely depends on host cell-derived membranes (Fig. 2G, S2D), that play a crucial role in animal cell entry and viral infectivity (Fig. 2H).

Lastly, our findings have significant implications for deltavirus co-infections in humans. Although the evidence for extra-hepatic deltavirus presence in human tissues is scarce—limited to reports in minor salivary glands ^39,40^—our data underscores the need for systematic monitoring across diverse human tissues and organs. The ability of animal deltaviruses to be co-packaged with *Herpesviridae* and infect human neurons (Fig. 3H-I) raises the possibility of overlooked infections in the peripheral or central nervous systems (CNS). Given that several billion people live with HSV-1 ^41^ and that HSV-1 establishes latency in the nervous system ^42^, chronic deltavirus co-infection could exacerbate disease outcomes such as for Herpes Simplex Encephalitis ^43^ or age-associated neurodegenerative diseases ^42,44^. Along this line, SDeV was first identified in a boa constrictor afflicted with Boid Inclusion Body Disease (BIBD), a condition marked by severe CNS symptoms ^4,32^, further supporting a potential neurotropic impact of deltaviruses.

In conclusion, we identify a novel mode of viral spread among animal viruses, facilitated by a viral Trojan Horse, and reveal associations between deltaviruses and diverse viral families. These findings provide mechanistic insights into the host- and tropism-shifting capabilities of these circular RNA viruses.

## Acknowledgments

We would like to thank Sébastien Pfeffer (Institut de Biologie Moléculaire et Cellulaire, Strasbourg, France) for providing VSV-GFP virus, Nadine Laguette (Institut de Génétique Moléculaire de Montpellier, France) for providing HSV-1 KOS-64-GFP virus and Gert Zimmer (Institute of Virology and Immunology, Switzerland) for providing G-pseudotyped VSV∗ΔG-GFP virus. We thank Koichi Watashi (National Institute of Infectious Diseases, Japan) for providing the mmDeV construct and Dieter Glebe (University of Giessen, Germany) for providing the RDeV construct. We thank the laboratory of Marcel Leist (Konstanz University, Germany) for the gift of LUHMES cells. We thank Dr. Eric Kremmer (Institut de Génétique Moléculaire de Montpellier, France) for providing the Citrine-encoding plasmid and Pr. Richard M Longnecker (Northwestern University, USA) for providing HSV-1 gD, gB, gH and gL plasmids. We thank Aurélie Ancelin (Centre de Biologie Structurale, France) for piloting the cryo-EM microscope. We thank the CEMIPAI (UAR 3725 CNRS/Montpellier University) BSL3 Facility for access to their facilities and thank their personnel. Finally, the authors acknowledge the imaging facility MRI, a member of the national infrastructure France-BioImaging supported by the French National Research Agency (ANR-10-INBS-04).

## Author contributions

Conceptualization: JM, KM

Methodology: JM, AF, AT, SDR, MPB, SL, FS, HDR, SD, KM

Investigation: JM, AF, FS, SL, RS, JH, LK Visualization: JM, AF, SL, MPB, FS

Funding acquisition: KM

Supervision: JM, KM

Writing – original draft: JM, KM

Writing – review & editing: JM, AF, SL, SG, RG, JH, KM

## Data and materials availability

All data is available upon request to the corresponding author.

## Supplemental information

Figures S1-4: Supporting information for main results.

Movie S1-3: 3D renditions of panels from Figure 3 and S3.

## Supplementary Materials for

### Materials and Methods

#### Cell lines

NIH3T3 (CVCL_0594), Vero (CVCL_0059) and BHK-21 (CVCL_1914) were grown in Dulbecco’s Modified Eagle Medium (DMEM), high glucose (Thermo Fisher Scientific) supplemented with 10% Fetal Bovine Serum (FBS) (Dutscher) and Antibiotic-Antimycotic 1X (Thermo Fisher Scientific) at 37°C, 5% CO_2_. I/1Ki (CVCL_B6ZK) were maintained in MEM, low glucose (ThermoFisher Scientific), supplemented with 10% FBS (Gibco), 2 mM L-glutamine (ThermoFisher Scientific), 100 IU/ml penicillin (ThermoFisher Scientific) and 100 ug/ml streptomycin (ThermoFisher Scientific) at 30°C, 5% CO2. NIH3T3-RDeV were generated previously ^1^ and NIH3T3-mmDeV were generated likewise. I/1Ki-SDeV were established by transfecting I/1Ki cells with 1.2xSwSCV-1-FWD plasmid as described in ^2^. LUHMES cells were cultured and differentiated as previously described (Kozoriz et al. in press). Briefly, LUHMES cells were maintained at 37°C in a humidified 5% CO2 atmosphere, in culture dishes previously coated with 50 mg/ml poly-L-ornithine and 1 mg/ml fibronectin, with the addition of 10 mg/ml laminin for glass coverslips. Proliferation medium comprised Advanced DMEM/F12 medium supplemented with N2 supplement, 2 mM GlutaMAX, 40 ng/ml bFGF and 100 IU/ml-100 µg/ml penicillin-streptomycin. For differentiation into dopaminergic neurons, LUHMES cells were switched to differentiation medium (Advanced DMEM/F12 medium supplemented with N2 supplement, 2 mM GlutaMAX, 1 mM dbcAMP, 1 µg /ml tetracycline, 2 ng/ml GDNF and 100 IU/ml-100 µg/ml penicillin-streptomycin) for 48 hours and then seeded at a density of 128,000 cells/cm^2^ in differentiation medium on glass coverslips. LUHMES-derived neurons were infected after 5 days of differentiation.

#### Plasmids and transfections

The VSV-G encoding plasmid (pMD.G) was described previously ^3^. HSV-1 glycoprotein (UL27 (gB), UL1 (gL), UL22 (gH) and US6 (gD)) coding plasmids were a gift from Prof. Richard M Longnecker (Northwestern University, USA) ^4^. HCMV glycoproteins (gH, gL, gO) coding sequences were synthesized as gBlocks (IDT) and cloned into pcDNA3.1 backbones by Gibson Assembly (New England Biolabs). The mCitrine-encoding plasmid was kindly provided by Dr. E Kremer. Plasmids were controlled by Sanger sequencing (Eurofins) or whole plasmid sequencing (Eurofins), when necessary. Transfections of plasmids were performed by reverse transfection of NIH3T3-RDeV cells in 24-well plates using the JetPei reagent (Polyplus), following the manufacturer’s instructions.

#### Viral stock productions and titrations

For VSV-GFP production, BHK-21 were seeded into 25 cm^2^ flasks and grown to 80% confluency and inoculated with 1 μl stock viral particles gifted by Dr Sébastien Pfeffer (Institut de Biologie Moléculaire et Cellulaire, France) in DMEM without FBS and left overnight at 37°C. Sixteen hours post-infection, the cell culture medium was harvested, cleared by centrifugation in a Heraeus Multifuge X3 FR (Thermo Fisher Scientific) using a TX-1000 swinging rotor (20 min, 3000 rpm, 4°C), aliquoted and stored at -80°C. Titration was performed on NIH3T3 cells by serial dilution infection for 1 h followed by a PBS wash and incubation in complete DMEM for 8 h. Quantification of the percentage of GFP-positive cells was performed following fixation (4% paraformaldehyde (PFA), 20 min, room temperature (RT)) and DAPI staining, using an ImageXpress Pico microscope (Molecular Devices) with a 4x lens.

G-pseudotyped VSV∗ΔG-GFP (VSVΔG-GFP+G) particles were a gift from Dr. Gert Zimmer (Institute of Virology and Immunology, Switzerland) ^5^. For amplification, BHK-21 cells were transfected with a VSV-G expressing plasmid. 24 hours after, they were infected with VSVΔG-GFP+G at MOI 0.1 for 1 h at 37°C, followed by a PBS 1x wash and resuspension in DMEM without serum. 16 hours later viral supernatants were recovered and following steps were carried out as above for VSV-GFP.

HSV-1 KOS-64-GFP strain (HSV-1-GFP) particles were a gift from Dr. Nadine Laguette (Institut de Génétique Moléculaire de Montpellier, France). Two 175 cm^2^ flasks of Vero cells were infected at MOI 0.1 for 1 h in DMEM without FBS, washed in PBS 1x, and left for 72 h in DMEM supplemented with 1% FBS at 37°C. The viral supernatant was cleared by centrifugation in a Heraeus Multifuge X3 FR (Thermo Fisher Scientific) using a TX-1000 swinging rotor (1 h, 3000 rpm, 4°C) and low speed ultracentrifuged (4 h, 12500 rpm, 4°C) in an OPTIMA L-80 XP (Beckman) ultracentrifuge using a SW 32Ti swinging rotor. The pellet was rehydrated in 200 μl DMEM without FBS for 1 h at 4°C, resuspended, aliquoted and stored at -80°C. Titration was performed on Vero cells by serial dilution infection for 1 h followed by a PBS 1X wash and incubation in complete DMEM for 48 hours and quantification of the number of GFP-positive cells following fixation (4% PFA, 20 min, RT) and DAPI staining, using an ImageXpress Pico microscope (Molecular Devices) with a 4x lens.

The UGV-1 viral stock utilized in this study was produced as described in ^6^.

#### Infections, superinfections and virus spotting

For VSV-GFP or VSVΔG-GFP+G infections of NIH3T3-WT or superinfections of NIH3T3-RDeV or NIH3T3-mmDeV cells, 80% confluent 75 cm^2^ flasks were infected at MOI 0.5 for 1 h in DMEM without FBS, washed once with PBS 1x and incubated for 16 hours in DMEM without FBS at 37°C. Viral supernatants were cleared by centrifugation in a swinging rotor (20 min, 3000 rpm, 4°C). Low Speed Centrifugation (LSC) was performed when necessary as descrived in ^7^. LSC consisted in centrifugation (18 hours, 5300xg, 15°C) using a Heraeus Multifuge X3 FR (Thermo Fisher Scientific) using a Fiberlite F15-8 x 50cy fixed angle rotor, followed by resuspension of LSC pellet in 100-150 μl DMEM without FBS. For incoming virus experiments, Vero cells were seeded at low confluency onto coverslips in 24-well plates and left to attach and spread for 6 hours. Cells were incubated with 1X CellTrace-CFSE (Thermo Fisher Scientific) in DMEM without FBS for 20 min at 37°C, washed twice with DMEM 10% FBS and incubated in DMEM 10% FBS for 10 min at 37°C. Cells were washed with PBS 1X and infected with VSV-RDeV Input, SNT or PEL fractions for 30 minutes at 37°C. Cells were then washed with PBS 1x and fixed (4% PFA, 20 min, RT). Cells were washed twice with PBS 1X and processed for immunofluorescence.

For HSV-1-GFP infections of NIH3T3-WT or superinfections of NIH3T3-RDeV cells, two 80% confluent 175 cm^2^ flasks were infected at MOI 0.1 for 1 hour in DMEM without FBS, washed in PBS 1x and left for 72 hours in DMEM 1% FBS. The viral supernatant was cleared by centrifugation in a Heraeus Multifuge X3 FR (Thermo Fisher Scientific) using a TX-1000 swinging rotor (1 hour, 3000 rpm, 4°C) and low speed ultracentrifuged (4 hours, 12500 rpm, 4°C) in an OPTIMA L-80 XP (Beckman) ultracentrifuge using a SW 32Ti swinging rotor. The pellet was rehydrated in 200 μl DMEM without FBS for 1 hour at 4°C then resuspended in DMEM without FBS. SNT and PEL fraction infections of Vero cells were performed for 1 hour at 37°C in DMEM without FBS, washed once with PBS 1x and fresh complete DMEM supplemented with either 50 μM Acyclovir (APExBIO) or DMSO. Seventy-two hours later cells were fixed (4% PFA, 20 min, RT), washed thrice in PBS 1x and processed for immunofluorescence. For Input, SNT and PEL fraction infections of LUHMES-derived neurons, infections were performed for 1 hour at 37°C in differentiation medium and replaced with fresh differentiation medium supplemented with 50 μM Acyclovir (APExBIO) or DMSO. Cells were lysed 2- or 3-days post-infection and processed for RT-qPCR or were fixed 3-days post-infection (4% PFA, 20 min, RT), washed once in PBS 1x and processed for immunofluorescence.

For UGV-1 infections of I/1Ki-WT or superinfections of I/1Ki-SDeV, cells were inoculated with UGV-1 at MOI 0.1 and following 1 hour adsorption at 30°C, the inoculum was replaced with fresh fully supplemented DMEM. The media was replaced at 4- and 7-days post-infection with fresh fully supplemented DMEM. At 9 days post-infection (dpi), cells were washed twice with a 1:1 mix of Freestyle 293:Freestyle CHO (Thermo Fisher Scientific) supplemented with 2 mM L-glutamine, and kept in this medium until the supernatants were collected at 12 dpi. Samples were cleared (10 min, 5000xg, RT) in a Fresco 21 centrifuge (Thermo Fisher Scientific) and concentrated in Amicon-Ultra centrifugal filters (Merck) in a SL 16R centrifuge (Thermo Fisher Scientific). This input was then centrifuged (4 h, 20000 xg, RT) in a Fresco 21 centrifuge and the pellet was resuspended in 150 μl of PBS 1x. Calculation of the infectivity/ml of SDeV SNT and PEL fractions was performed as previously described ^8^. Briefly, a 10-fold dilution series of the SNT and PEL fractions was performed. The diluted virus was dispensed onto I/1Ki-WT cells grown on collagen-coated ViewPlate black 96-well plates at 100 μl per well. Four days post-infection, cells were fixed (4% PFA, 20 min, RT) and stained for snake delta Antigen (SDAg) and DAPI. The number of SDAg-expressing cells were quantified per condition, and Focus Forming Units (FFU)/ml of each fraction was determined.

For virus spotting experiments, 12 mm coverslips were pre-coated with 0.01% Poly-L-Lysin for 1 hour in 24-well plates, washed once in PBS 1x and virus fractions were spotted onto them for 5 min at RT, followed by dilution into 500 µl final volume PBS 1x. Coverslips were rested for up to 72 hours at 4°C and then fixed (4% PFA, 20 min, RT), washed thrice with PBS 1x and then processed for immunofluorescence.

For glycoprotein-pseudotyped RDeV infection experiments, Vero cells were seeded into 96-well plates at low density, infected for 1 hour, washed with PBS 1X, replaced with complete DMEM and left 48 hours at 37°C. Cells were then fixed (4% PFA, 20 min, RT) and washed thrice with PBS 1x and processed for immunofluorescence. Deltavirus- and helper virus-containing stocks were directly processed for experiments and any excess were discarded.

#### Immunoblotting

Viral supernatants were supplemented with Laemmli buffer (Bio-Rad), 10% β-mercaptoethanol (Bio-Rad) and heated at 95°C for 10 min. Samples were loaded onto 4–20% Mini-PROTEAN TGX gels (Bio-Rad) and SDS-PAGE electrophoresis was performed at 110 V until desired migration. Proteins were transferred onto an ethanol-activated PVDF membrane (Bio-Rad) for 7 min, 2.5 A, 12 V using a Trans-Blot Turbo Transfer System (Bio-Rad). Blocking was performed in PBS 1x-5% milk for 1 h at RT. Primary antibodies (Human Ig-Patient1 for RDAg, 1:1000, ^1^; Mouse anti-VSV-M; 1:500, Kerafast; Mouse anti-HSV-1-ICP0, 1:100, Santa Cruz) were incubated overnight in PBS 1x-5% milk at 4°C in rotation. Membranes were washed thrice in PBS 1x-0.1% Tween 20 (Bio-Rad) (PBST) for 5 min. Secondary antibodies (anti-Human-HRP, 1:2500; anti-mouse-HRP, 1:2000; Goat anti-human-IRDye800, 1:2500, LI-COR, Donkey anti-mouse-IRDye680, 1:2500, LI-COR) were incubated for 1 h at RT in PBST and washed thrice in PBST. Revelation of membranes was performed either directly using an Odyssey M Infrared Imaging System (LI-COR Biosciences) or following Clarity ECL Western Blotting Substrate (Bio-Rad), by incubation for 5 min at RT using a ChemiDoc system (Bio-Rad). Images were post-processed using FIJI ^9^. SDeV and UGV-1 supernatant immunoblots were performed as previously described ^10,8^.

#### Reverse Transcription quantitative Polymerase Chain Reaction (RT-qPCR)

In the case of virus supernatants, samples were lysed using the Power SYBR Green Cell-to-CT Kit (Invitrogen). The RT and qPCR were performed following the manufacturer’s instructions using a CFX Opus 384 Real-Time PCR System (Bio-Rad). Primers used in this study are as follows: RDeV genome: FWD 5’-CAAGAAAAACCACACTGCACA-3’, REV 5’-AGGGCAAAGAGAACAGACGA-3’; VSV (L): FWD 5’-CATGATCCTGCTCTTCGTCA-3’, REV 5’-TGCAAGCCCGGTATCTTATC-3’; SDeV genome: FWD 5′-GAAAGACGCGACAACTGTGAGTC-3′, REV 5′-GTCTAGTCCCGTTCCGGTTCTATG-3′, and probe 5′ 6-Fam (carboxyfluorescein)-GGAGATCCGAGAGGGGAGAAGAGGAGAGGTC-BHQ (black hole quencher)-1 3′; UGV-1 (S): FWD 5′-CAAGAAAAACCACACTGCACA-3, REV 5′-AACCTGTTGTGTTCAGTAGT-3′, and probe 5′-6-Fam-CTCGACAAGCGTGGGCGGAGG-BHQ-1-3′. Data were processed using the CFX manager suite (Bio-Rad). For viral supernatant analysis final values were normalized based on total volume of viral fractions. For LUHMES-derived neuron experiments, cells were washed once with PBS 1x and processed as mentioned above. Values were normalized to cellular ribosomal RNA 18S levels. SDeV and UGV-1 supernatant RNA extractions and qRT-PCRs were performed as previously described ^8,11^.

#### Immunofluorescence, epifluorescence, confocal, Airyscan and STED microscopy

For immunofluorescence processing, viruses or cells were permeabilized in 0.2% Triton X-100 in PBS 1x for 10 min at RT, followed by quenching and blocking in NGB buffer [50 mM NH4Cl, 2% Goat serum, 2% Bovine serum albumin, in PBS 1x] for 1 h in the dark. Primary antibodies (Human Ig-Patient1 for RDAg, 1:1000, ^1^; Mouse anti-VSV-M; 1:500, Kerafast; Mouse anti-VSV-G; 1:300, Kerafast; Mouse anti-HSV-1-VP5, 1:250, Santa Cruz; Mouse-anti-HSV-1-ICP0, 1:50, Santa Cruz; Rabbit anti-SDAg; 1:1000, ^12^; Rabbit anti-UHV-NP-coupled Alexa Fluor 594; 1:500, ^8^; Mouse anti-Tuj1, 1:500, BioLegend) were diluted in NGB and incubated for 1 h at RT in the dark. Coverslips were washed thrice in PBS 1x and incubated with secondary antibodies (For confocal: Goat anti-human Alexa Fluor 647, 1:1000, Thermo Fisher Scientific; Donkey anti-human Alexa Fluor 555, 1:1000, Thermo Fisher Scientific; Donkey anti-mouse Alexa Fluor 555: 1:1000, Thermo Fisher Scientific; Goat anti-rabbit Alexa Fluor 488, 1:1000, Thermo Fisher Scientific; For STED: Donkey anti-human-STAR-RED, 1:100, Abberior; Goat anti-Rabbit-STAR-RED, 1:100, Abberior; Goat anti-mouse Alexa Fluor 594, 1:100, Thermo Fisher Scientific) for 1 h at RT in the dark and washed thrice with PBS 1x. In the case of LUHMES, secondary antibody mix was supplemented with Phalloidin Alexa Fluor 647 (1:100, Thermo Fisher Scientific), when necessary. If needed, cells were incubated with DAPI (Thermo Fisher Scientific) to visualize nuclei. Coverslips were then washed once in milliQ water. For confocal imaging, coverslips were mounted onto glass slides using Prolong Gold Antifade Mountant (Thermo Fisher Scientific) and for STED imaging, mounted in solid antifade mounting medium (Abberior GmbH). For glycoprotein-pseudotyped RDeV infection experiments, cells were stained for RDAg and DAPI as above and imaged directly in the 96-well plates using an ImageXpress Pico microscope (Molecular Devices) with a 4× air lens.

Confocal images were acquired on an LSM980 8Y with Airyscan 2 module (ZEISS) using a 63× oil immersion objective. Pre-processing of raw Airyscan images was performed on the ZEN Blue software (ZEISS). Postprocessing of all images was performed using the FIJI software. STED images were acquired on an ExoertLine STED microscope (Abberior Instruments GmbH) using a 100× oil immersion objective with N.A. 1.4. STAR-RED and Alexa Fluor 594 were excited at 640nm and 561nm respectively, and depleted by a 775nm STED laser. Collection of fluorescence emissions were set as follows: 650-750nm for STAR-RED, 570-630nm for Alexa Fluor 594. For incoming virus experiments, CellTrace-CFSE was used as a third staining to provide intracellular context to STED imaging of viral particles. Imaging of CellTrace-CFSE was done in confocal mode, sequentially with STED imaging. Excitation was done at 485nm and fluorescence emission was collected on the 500-550nm range. Postprocessing of images was performed using the FIJI software.

#### Negative Stain, Immunogold- and Transmission-Electron Microscopy

For NS-TEM, 3 µl of viral fractions were applied to glow-discharged grids and incubated 2 mi. Following this, treatment with 0.75 % uranyl acetate was done for an additional 2 min. Grids were loaded onto a 120 kV transmission electron microscope and imaged using the JEOL software. Images were collected images were then post-processed on FIJI. For Immunogold-EM, Nickel formvar carbon grids were incubated for 60 min with each sample. After 3 washes in wash buffer [0,1% BSA, PBS 1x], grids were incubated 10 min in permeabilization buffer [0,1% BSA, 0,2% Triton-X100, PBS 1x]. After 6 washes in wash buffer grids were incubated 20 min in BSA-c saturation solution. Following 4 washes in wash buffer, grids were incubated for 1 h in solution containing a 1:100 dilution of human anti-HDV polyclonal antibody, or a 1:100 dilution of mouse anti-VSV-G (8G5F11, Merck Millipore) or both. Following 6 washes in wash buffer, the grids were incubated in wash buffer supplemented with cognate gold-conjugated antibody (1:30, Aurion), goat anti-mouse 10 nm gold beads (1:30) or goat anti-human 6 nm golds beads (1:30). The grids were finally washed 6 times in wash buffer and 6 times in distilled water. Contrast staining was achieved by incubating the grids briefly in a 2% uranyl acetate diluted in water (1:2). The grids were observed by transmission with a JEM-1011 (JEOL) transmission electron microscope operated at 100 kV and equipped with an Ametek-GATAN RIO9 CMOS camera.

#### Cryo-Electron Microscopy

For Cryo-EM experiments, 3 µl of viral fractions were applied to glow-discharged (25 mA, 10 s) QuantiFoil Copper R 1.2/1.3 300-mesh holey carbon grids (QuantiFoil, Micro Tools GmbH, Germany), blotted for 4.5 s, and then flash-frozen in liquid ethane using a Leica EM GP2. Images were acquired using a JEOL2200FS microscope operating at 200 kV and equipped with an omega energy filter at the PIBBS platform localized at the CBS (Montpellier, France). Micrographs were recorded by hand on a GATAN K3 detector using the serialEM software. Micrographs were then post-processed on FIJI.

#### Atomic Force Microscopy

Glass coverslips were coated with 0.01% poly-l-lysine (Sigma) for 1 h and VSV-infected cell supernatants were spotted onto them, rested for 5 min and resuspended in PBS 1x and stored at 4°C for 48 h. The coverslips were mounted into a JPK CoverslipHolder (Bruker) filled with 1 mL PBS 1x. AFM imaging was performed in PBS medium at room temperature on a NanoWizard IVxp atomic force microscope (JPK BioAFM, Bruker Nano GmbH, Berlin, Germany) operating in a biosafety level 3 laboratory. Topographic images were obtained using the PeakForce Tapping mode with either BL-AC40TS probes (Olympus, nominal spring constant of 0.09 N/m) or qpBioAC-CB3 (Nanosensors, nominal spring constant 0.06 N/m). Before each set of acquisitions, the cantilever sensitivity and spring constant were calibrated (thermal noise method). The applied force was kept at 130 pN with a PeakForce frequency of 1.6kHz and amplitude of 70nm. Using the SPM-data processing software (Bruker Nano GmbH, Berlin, Germany) images were flattened with a polynomial/histogram line fit. Low-pass Gaussian and/or median filtering was applied to remove minor noise from the images. The Z-color scale in all images is given as relative after processing. Particle height analysis was performed using the cross-section tool of the analysis software to calculate the maximal central height of each particle.

#### Image analysis

For quantification of NS-TEM images: 1-for free deltavirus or VSV-deltavirus particle number and size, images were manually screened and particles identified using FIJI. Particles were isolated using the freehand selection tool, particle sizes were assessed and exported using the measure tool. From the surface area, was approximated the diameter using ø=2√(A/π). 2-For UGV-1-SDeV particle quantifications, UGV-1 particles were manually identified using FIJI and the presence or not of intra-virion densities within the UGV-1 particles was assessed.

STED deconvolution was performed for VSV and UGV-1 images with Huygens Professional 24.10 software (Scientific Volume Imaging B.V.) using the classic maximum likelihood algorithm (CMLE). Automatic background estimation was based on the lowest mode, signal to noise (SNR) ratio was between 10 and 30, and acuity parameters were between 0 and 70. For HSV-1 images STED deconvolution was performed with the TRUESHARP image boosting tool (Abberior; https://app.truesharp.rocks/) using the following settings: Number of iterations: 5, PSF size: 2, Background size: 8 and Background weight: 1.

For quantification of STED images: 1-for VSV images, segmentation of VSV and deltavirus particles was performed with custom made Python™ routines. Briefly, a blob detection algorithm (based on a Laplacian filter) was employed to detect punctate deltaviruses and subsequent segmentation of deltavirus particles was performed using a defined segmentation threshold. This threshold was rescaled by the statistical properties of the filtered image in order to determine a common threshold value between repeats. VSV segmentation was performed with Cellpose ^13^, given that its bullet shaped virion is more complex to detect. Following this, any VSV aggregates were removed as the number of VSV particles in these aggregates was not definable and the corresponding projection onto the deltavirus images were also removed. In some cases, where an aggregate was not removed by this process, it was removed manually. Following segmentation, extraction of the of the position of the center of mass of each segmented deltavirus and projected onto the VSV segmentation. In the case of a non-overlapping deltavirus, we expanded the segmentation recursively until an overlapping occurred. The overlapping distance between the center of mass of each deltavirus and a VSV was recorded. A maximum distance between the deltavirus center of mass and the edge of a VSV virion of 2 pixels (40 nm) was considered overlapping, and the rest were not. 2-For HSV-1 images, FIJI was used to detect HSV-1 particles by thresholding the HSV-1 channel. The analyze particles tool was then used to isolate HSV-1 particles. Identified particles were then screened for presence or absence of RDAg. 3-For UGV-1 images, analysis was performed as for VSV, but the UGV-1 particles were not detected with Cellpose, but with a blob detection algorithm (based on Difference of Gaussian), as particles were circular in shape.

For quantifications of the immunogold images: VSV particles were identified by 10 nm gold beads and the presence or absence of 6 nm gold beads was noted in both VSV-WT and VSV-RDeV conditions. The percentage of 6 nm bead-positive VSV particles was then manually counted and the percentage of positivity was inferred.

#### Statistical analysis

For all statistical tests used, the specific test is indicated in the figure legends. All statistical analysis was performed using the GraphPad PRISM software. ns: non-significant, *: p<0.05, **: p< 0.01, ***: p<0.001, ****: p<0.0001.

**Fig. S1.**
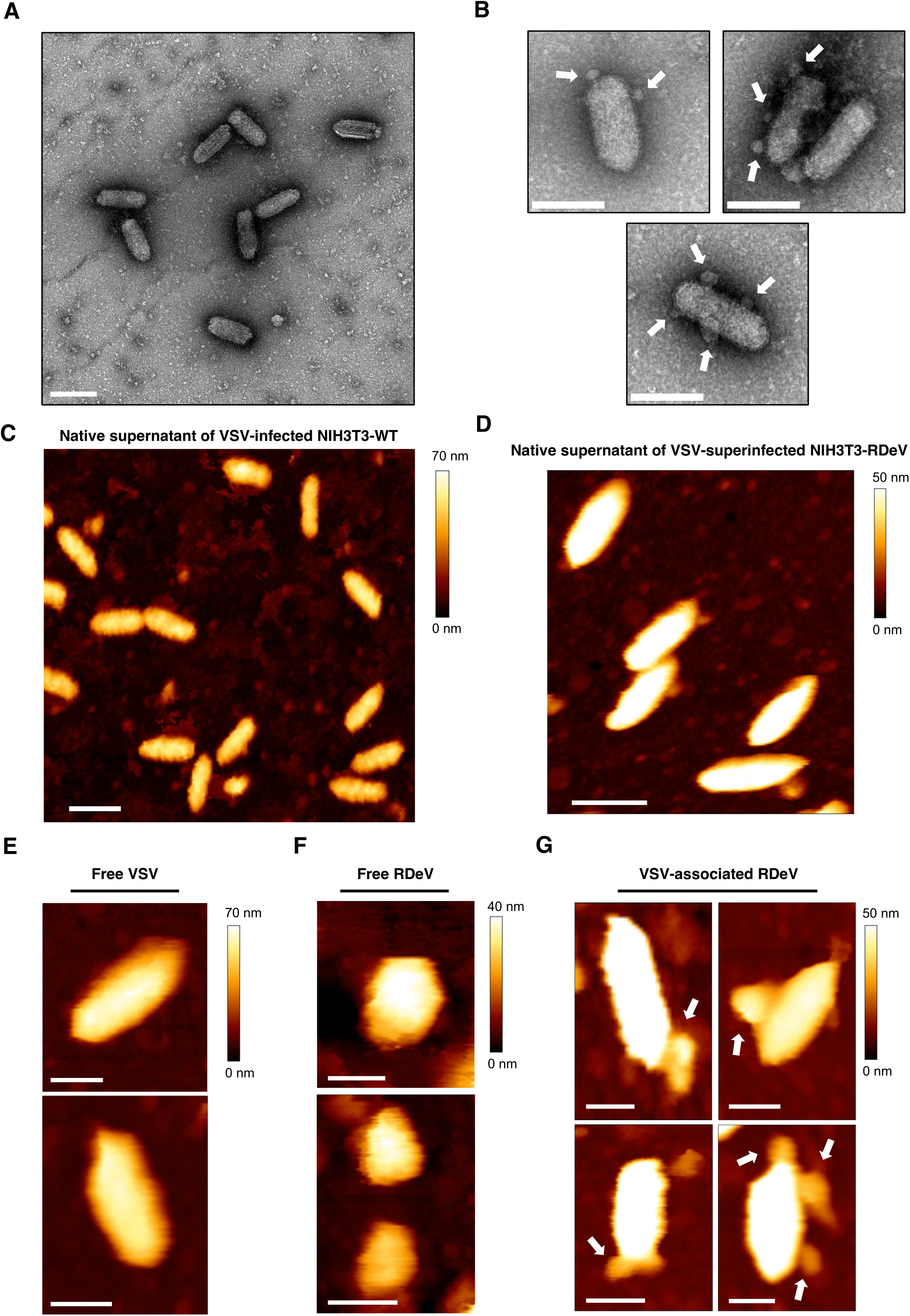
(**A**) NS-TEM image of VSV-infected NIH3T3-WT supernatants. Scale bar: 200 nm. (**B**) NS-TEM images of VSV virions with multiple RDeV-associated particles. Scale bar: 200 nm. (**C**) Representative field of native VSV particles from NIH3T3-WT supernatants by AFM. Scale bar: 200 nm. (**D**) AFM image of non-fixed VSV-superinfected NIH3T3-RDeV cell supernatants. Scale bar: 200 nm. (**E-G**) AFM images of single particles of non-fixed VSV (**E**), free RDeV (**F**) and VSV-RDeV (**G**) from VSV-superinfected NIH3T3-RDeV cell supernatants. Scale bars: E-100 nm, F-50 nm, G-100 nm. Experiments are representative of 2 independent repeats.

**Fig. S2.**
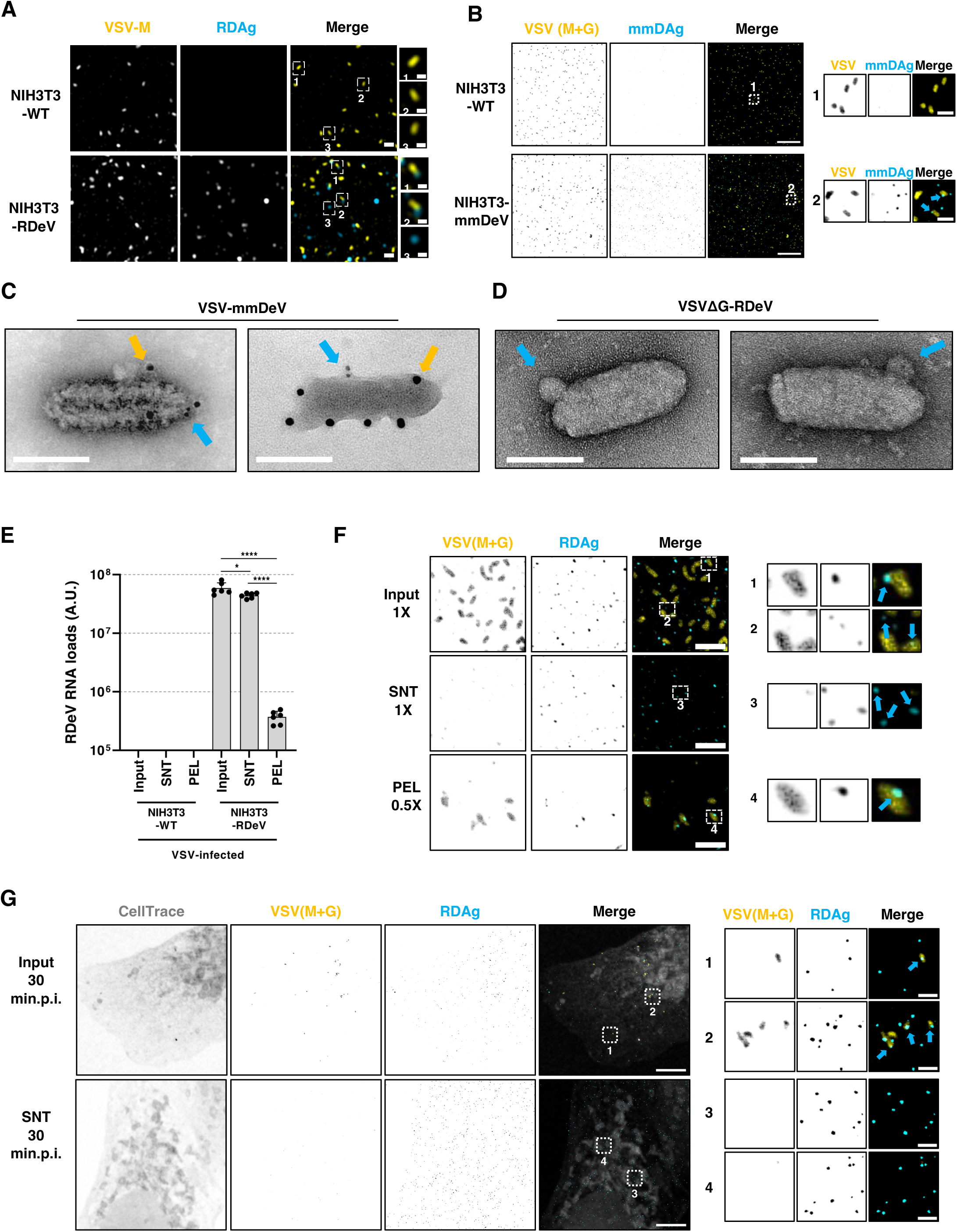
(**A**) Representative Airyscan images of VSV-infected NIH3T3-WT or -superinfected NIH3T3-RDeV supernatants. VSV (M+G) is represented in yellow and RDAg in cyan. Single channels are shown in grayscale. Scale bar: 500 µm, inset: 200 nm. (**B**) Representative STED images of VSV-infected NIH3T3-WT or -superinfected NIH3T3-mmDeV supernatants. VSV (M+G) is represented in yellow and *Marmota monax* delta-antigen (mmDAg) in cyan. Single channels are shown in inverted grey. Cyan arrow points to VSV-mmDeV particles. Scale bar: 5 µm, inset: 500 nm. (**C**) Representative NS-TEM images of immunogold labelled VSV-M (10 nm gold beads; yellow arrows) and mmDAg (6 nm gold beads; cyan arrows) from supernatants of VSV-superinfected NIH3T3-mmDeV. Scale bar: 100 nm. (**D**) NS-TEM images of VSVΔG-RDeV virions produced from VSVΔG+G-superinfected NIH3T3-RDeV 16 h post-infection. Scale bar: 100 nm. (**E**) RDeV RNA RT-qPCR of volume equilibrated LSC fractions of 16 h VSV-infected NIH3T3-WT or NIH3T3-RDeV supernatants. (**F**) Representative STED images of VSV-RDeV LSC 1X input, 1X SNT and 0.5X PEL fractions (PEL for pellet fraction and SNT for supernatant fraction). Insets show free RDeV and VSV-RDeV particles in the different fractions. Scale bar: 1 µm (**G**) STED image of Vero cells prestained with Celltrace-CFSE and infected with VSV-RDeV input or LSC SNT fraction for 30 min. VSV (M+G) are stained in yellow and RDAg in cyan. Cyan arrows point to VSV-RDeV particles among intracellular membranes. Scale bar: 5 µm: inset: 500 nm. Experiments are representative of 2 (A-D, F, G) or 3 (E) independent repeats, performed in technical duplicates for E. Unpaired T-tests were used to evaluate statistical significance.

**Fig. S3.**
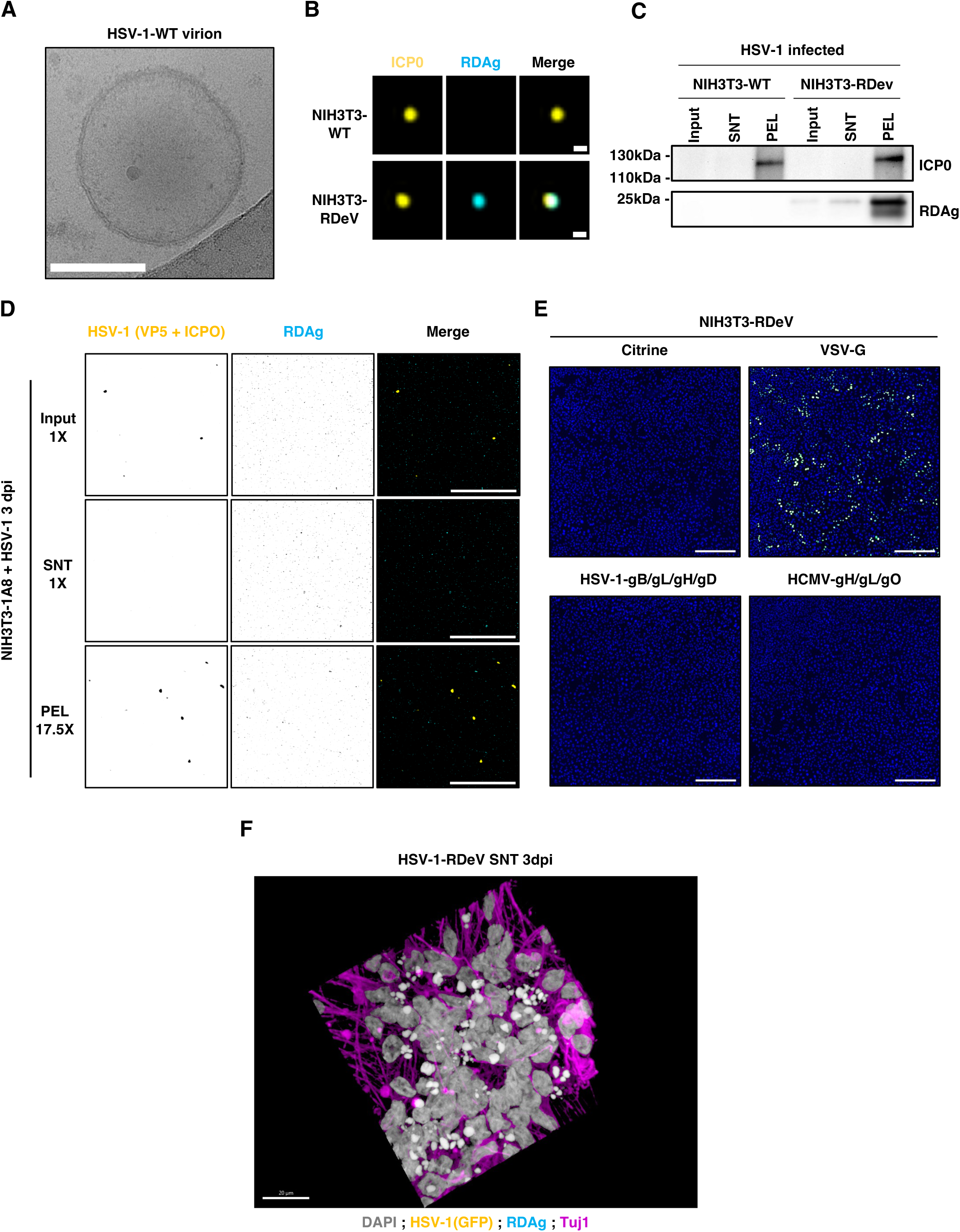
(**A**) Example of an HSV-1 virion from HSV-1-infected NIH3T3-WT cells 72 h post-infection, observed by our Cryo-EM approach. Scale bar: 100 nm. (**B**) Representative Airyscan image of individual HSV-1 virions from HSV-1-infected NIH3T3-WT or -superinfected NIH3T3-RDeV supernatants. HSV-1 (ICP0) is represented in yellow and RDAg in cyan. Scale bar: 200 nm. (**C**) Immunoblot using equal volumes of low-speed ultracentrifugation Input (1X), SNT (1X) and PEL (250X) fractions from HSV-1-infected NIH3T3-WT or -superinfected NIH3T3-RDeV supernatants (PEL for pellet fraction and SNT for supernatant fraction). (**D**) Representative STED images of HSV-1-RDeV low-speed ultracentrifugation input (1X), SNT (1X) and PEL (17.5X) fractions. Scale bar: 5 µm. (**E**) Representative fields of Vero cells 3 days post-infection, infected with supernatants from NIH3T3-RDeV transfected with either Citrine, VSV-G, HSV-1-gB/gL/gH/gD or HCMV-gH/gL/gO encoding plasmids for 2 days. RDAg is shown in yellow and DAPI in blue. Scale bar: 500 µm. (**F**) Representative confocal z-stack 3D-rendering of a field of LUHMES-derived neurons infected with HSV-1-RDeV SNT fraction 3 days post-infection in the absence of Acyclovir. Nuclei are shown in grey, HSV-1 (GFP) in yellow, RDAg in cyan and Tuj1(neuronal marker) in magenta. Scale bar: 20 µm. Corresponding animated 3D-rendering of this z-stack can be found in Movie S3. Experiments are representative of 2 (A-D) or 3 (E, F) independent repeats. Unpaired T-tests were used to evaluate statistical significance.

**Fig. S4.**
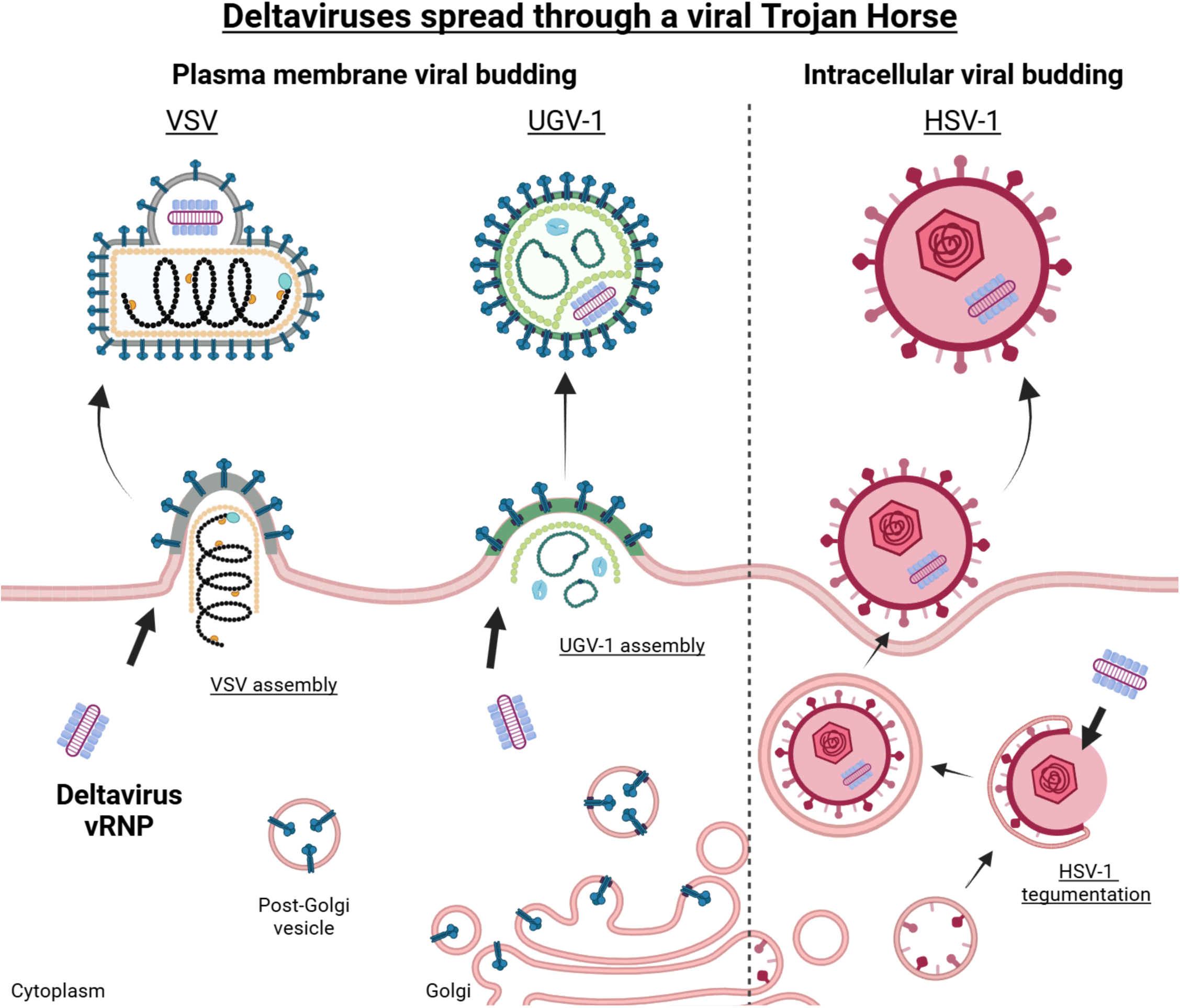
Schematic representation of the formation of viral Trojan Horses containing deltavirus vRNPs in the case of VSV, UGV-1 and HSV-1.

**Movie S1.**

Confocal z-stack 3D-rendering of a field of LUHMES-derived neurons infected with HSV-1-RDeV PEL fraction 3 days post-infection in the absence Acyclovir. Nuclei are shown in grey, HSV-1 (GFP) in yellow, RDAg in cyan and Tuj1 in magenta. Scale bar: 20 µm. Corresponding figure: Fig. 3H.

**Movie S2.**

Airyscan z-stack 3D-rendering of a field of LUHMES-derived neurons infected with HSV-1-RDeV PEL fraction 3 days post-infection in the absence Acyclovir. Nuclei are shown in grey, HSV-1 (GFP) in yellow, RDAg in cyan and Tuj1 in magenta. Single channels are shown in inverted grays. Scale bar: 10 µm. Corresponding figure: Fig. 3I.

**Movie S3.**

Confocal z-stack 3D-rendering of a field of LUHMES-derived neurons infected with HSV-1-RDeV SNT fraction 3 days post-infection in the absence Acyclovir. Nuclei are shown in grey, HSV-1 (GFP) in yellow, RDAg in cyan and Tuj1 in magenta. Scale bar: 20 µm. Corresponding figure: Fig. S3F.

